# Boosting ascomycin production through n-butanol addition in the fermentation process of *Streptomyces hygroscopicus var. ascomyceticus* ATCC 14891

**DOI:** 10.1101/2023.10.07.561346

**Authors:** Jinyu Meng, Qi Chen, Youyuan Li, Shuhong Gao, Daojing Zhang

## Abstract

Ascomycin (FK520) is a macrolide antibiotic known for its immunosuppressive activities. In this study, we screened several short-chain alcohols to enhance titer of FK520 in *Streptomyces hygroscopicus var. ascomyceticus* ATCC 14891, with a particular focus on n-butanol addition. After optimizing the n-butanol addition process, we achieved a FK520 yield of 569.37 mg/L with 0.8% n-butanol addition at 27 h, representing a 1.72-fold increase compared with the control group. We subsequently found that ROS levels of n-butanol addition group reached 1.49x10^5^ RFU/g biomass at 29 h, which was 3.02 times higher than the control group. The 0.8% n-butanol addition also promote the accumulation of biosynthesis precursors for FK520 production. The highest ethylmalonyl-CoA content surged to 93.1 nmol/g DCW at 48 h, marking a 5.3-fold increase. Likewise, the highest methylmalonyl-CoA and malonyl-CoA levels increased remarkable 4.33-fold and 3.33-fold compared with the control group at 72 h and at 120 h, respectively. Then, we explored the effects of oxygen supply on FK520 production with n-butanol addition, and improving oxygen supply caused a significant increase of FK520 in shake flask fermentation. Our research has revealed that addition of short-chain alcohols can regulate carbon flux toward FK520 biosynthesis by supplementing various CoA-esters, including ethylmalonyl-CoA, methylmalonyl-CoA, and malonyl-CoA.

## 1. Introduction

FK520, produced by various Streptomyces species such as *S. hygroscopicus* KK317 or *Streptomyces* sp. KCCM 11116p [1,2], is a nature 23-membered macrocyclic antibiotic with immunosuppressive properties [3], and effectiveness against malaria [4]. Furthermore, it displays antifungal and antispasmodic activities [5]. FK520 is highly regarded as a versatile and valuable drug [6]. Notably, as a structural analog of FK520, FK506 also is a nature 23-membered macrocyclic antibiotic, which is used for the treatment of xenograft rejection [7]. Most importantly, it can be used as an important chemical intermediates for the production of Pimecrolimus approved by FDA in 2001.

Pimecrolimus being semi-synthetic products of the Streptomyces-produced FK520, many researchers utilized *S. hygroscopicus var. Ascomyceticus* ATCC 14891 and its mutant strains to produce FK520. How to improve FK520 production has become hotspots in the world. Qi used femtosecond laser irradiation for strain mutagenesis, and mutant strain *S. hygroscopicus* SA68 was obtained. Then, with shikimic acid addition during fermentation of *S. hygroscopicus* SA68, FK520 titer was been improved to 270 mg/L [8]. Furthermore, they identified and engineered two potential gene manipulation targets (*pyc* and *fkbO*), resulting in an enhanced FK520 production of 610 mg/L [9]. Most of the studies about the enhancement of FK520 titer are associated with primary metabolic pathways inside of secondary metabolites for FK520 biosynthesis, which is regulated by precursor synthetic pathways and regulatory networks in *S. hygroscopicus var. ascomyceticus*. As we know, only particular signals such as stress from environmental could stimulate bacteria synthesize antibiotic [10]. And to discover the natural stimuli that activate antibiotic biosynthesis using novel methods has garnered significant interest among multiple research groups. Currently, substantial efforts are being dedicated to promoting the accumulation of antibiotics and stimulating the activity of silent gene clusters that regulate antibiotic biosynthesis using chemical elicitors, nonionic surfactants, and extracellular disturbances [11,12]. For instance, antibiotic biosynthesis in *S. antibioticus, S. griseus, and S. coelicolor* can be increased 2-25 fold by the stimulation of low concentrations of rare earth elements (REEs) [13]. Triton X-100, as a nonionic surfactant, can stimulate the biosynthesis of hypocrellin A in *Shiraia bambusicola* and its release with concentrations ranging from 0.5% to 3.0%[14]. As a chemical elicitor, DMSO can enhance the production of FK520 by 2.32-fold at a concentration of 0.6% in *S. hygroscopicus var. ascomyceticus* H16 [15].

As extracellular disturbances, short-chain alcohols could stimulate the overproduction of secondary metabolites in *Streptomyces* by causing environmental stress and altering the metabolic states of bacteria due to their toxicity [16,17], which has been reported by many research groups. For instance, isobutanol could promote the biosynthesis of AP-3 in *Actinosynnema pretiosum* [18]. n-propantol could improve the biosynthesis of erythromycin production in *Streptomyces erythreus* [19]. Ethanol could improve the validamycin A titer in *S.hygroscopicus* 5008 [20]. In these reports, apart from stimulating the biosynthesis of antibiotics, short-chain alcohols could also participate in the metabolic reactions of the bacteria by supplying the biosynthetic precursors of secondary metabolites. However, there have been no reports so far of promoting FK520 accumulation in *S. hygroscopicus var. ascomyceticus* ATCC 14891 through the short-chain alcohols supply.

In this research, we investigated the influence of various short-chain alcohols including n-propanol, isopropanol, n-butanol, isobutanol, 1, 3-propanediol on FK520 accumulation in *S. hygroscopicus var. ascomyceticus* ATCC 14891. Next, we explored the changes in fermentation process parameters and the variations in the content of CoA-esters to identify potential mechanisms that could regulate the carbon flux toward FK520 production. Simultaneously, we used the metabolic network model for simulating and analyzing how oxygen supply and n-butanol addition impact FK520 biosynthesis. Subsequently, based on the simulation and analysis results from GSMM, we further optimized the oxygen supply conditions by increasing the rotational speed during flask fermentation, thereby increasing the yield of FK520 with and without n-butanol addition. The insights gained from this research not only serve as a valuable reference but also provide guidance for future investigations into FK520 biosynthesis. Furthermore, these findings provide a rational metabolic engineering approach that can be applied to enhance other antibiotics accumulation with biosynthetic precursors similar to that of FK520, such as FK506 and Rapamycin.

## 2. Materials and methods

### 2.1. Strain and culture conditions

Throughout this study, the microorganism used was *S. hygroscopicus var. ascomyceticus* ATCC 14891. The components of the seed medium included the subsequent: 15g/L starch, 10g/L glucose, 10g/L soybean meal, 6g/L yeast extract, 2g/L NaCl, 1g/L (NH_4_)_2_SO_4_, 2g/L CaCO_3_, and a pH of 7.2. The fermentation medium in the shake flask included: 9g/L glycerol, 5g/L tryptone, 7g/L yeast extract, 2g/L corn steep liquor, 0.5g/L MnSO_4_, 1.5g/L (NH_4_)_2_SO_4_, 0.5g/L KH_2_PO_4_, 1g/L MgSO_4_·7H_2_O, 12g/L soybean oil, 1g/L CaCO_3_, with a pH of 6.7.

The growth of microorganism spores were conducted on Yeast Malt Extract Agar ATCC Medium 196 for 15 days at 28 °C. Afterward, the spores were introduced into 250 mL flasks with 25 mL of seed culture and incubated at 30 °C with agitation at 220 r/min for a duration of 44 hours.Following this, 2.5 mL of the seed culture medium was added to a 250 mL flask with 25 mL of fermentation culture. This medium was grown at 30 °C with agitation at 220 r/min and 250 r/min for a period of 5 days. All the short-chain alcohols, including n-propanol, isobutanol, isopropanol, 1,3-propanediol, and n-butanol, were acquired by sterile filtration through a 0.22 µm filter and were subsequently incorporated into the fermentation culture. The percentage (1.2%) short-chain alcohols (Fig. 1) were introduced into the batch medium after 24 h. Each group underwent three replicates.

### 2.2. Product, impurity, Glycerol, dry cell weight (DCW), and K_L_a determination

DCW was assessed by subjecting the cells to two washes with phosphate-buffered saline (PBS) after centrifugation at 8000 r/min for 10 minutes, followed by drying them at 105 °C until a consistent weight was reached.The ultimate biomass was quantified in grams per liter (g/L).

FK520 was extracted using methanol, and its production was determined by high-performance liquid chromatography (HPLC) according to the procedure outlined by Qi [21]. The HPLC system used was the 1260 Infinity II by Agilent Technologies, USA, equipped with a Zorbax SB-C18 analytical column (250 mm × 4.6 mm, Agilent Technologies, USA). In brief, 1 mL of the methanol was added with 1 mL of fermentation culture and then sonicated in water for 30 min. Following centrifugation at 12000 r/min for 5 min using a Centrifuge 5415 R (Eppendorf), the HPLC analysis was performed on the supernatant. The mobile phase used was a combination of acetonitrile and 0.1% phosphoric acid in water (in a proportion of 65:35, v/v) with a flow rate of 1 mL/min. A 10 μL sample introduction volume was used. The column temperature was kept at 55 °C, and the UV detector was configured to operate at 210 nm. The glycerol concentration was assessed through titration. [22]. K_L_a of different Rotational speed were assessed following the method outlined by Linek V[23].

The FK523 extraction determination was carried out in the same manner as for the product. Subsequently, the content of methanol-extracted sample was determined using ultra-high-performance liquid chromatography (UPLC, Ultimate 3000, Thermo Scientific) – electrospray ionization (ESI) – tandem mass spectrometry (MS, Q Exactive Plus, Thermo Scientific).The mobile phase was composed of two solvents, solvent A (50 mM ammonium formate solution) and solvent D (acetonitrile), at a flow rate of 300 µL/min, with the subsequent programmed: from 0.00 to 10.00 minutes, the ratio was 35% A and 65% D. The structure of impurity was identified using Full SCAN mode in positive mode.

### 2.3. Assessment of oxidative stress with n-butanol addition

The intracellular concentrations of reactive oxygen species (ROS) were assessed using the procedure outlined by Miranda, R. U [24].In brief, ROS-induced oxidation of the fluorogenic probe H2DCF-DA was monitored at various time during culture.5 mL of a 10 mM H2DCF-DA solution in cold phosphate-buffered saline (PBS) was added to 100 mg of mycelium. Then, the process was carried out under light-protected conditions at 37 °C for 30 min. Subsequently, each sample underwent three freeze-thaw cycles using liquid nitrogen. Following centrifugation at 12000 r/min for 5 minutes at 4 °C of each sample, 200 μL of the supernatant was moved to microplate reader (SYNERGY H1, BioTek, USA) with an excitation filter set at 488 nm and an emission filter set at 525 nm. The acquired signal was standardized based on biomass levels. Triplicate analysis was conducted for each sample.

### 2.4. Intracellular CoA-esters harvest and examination

Every sample underwent centrifugation at 12,000 r/min for 5 min to separate mycelium, followed by two washes with phosphate-buffered saline. CoA-ester extracts were prepared using a modified method based on the procedure described by Park, J. W [25]. The mycelium was reconstituted in 1 mL of 10% trichloroacetic acid (TCA). Following that, each sample was subjected to three rounds of freezing and thawing with liquid nitrogen and then underwent centrifugation at 12,000 r/min at 4 °C for 5 min. Each TCA sample was processed using an HLB Solid-Phase Extraction (SPE) cartridge from OASIS, following the specified protocol: 3 mL of methanol was initially introduced into the SPE cartridge, and then 3 mL of a 0.15% TCA solution was added to the cartridge. The TCA-extracted sample was loaded onto the cartridge, and a 1 mL water rinsing step followed. CoA-esters were eluted using 800 μL of methanol, and then the resulting eluate was dried at ambient temperature with Termovap Sample Concentrator. The dried samples were preserved in -80 °C for analysis. For the analysis, samples were dissolved in 80 µL of a 20 mmol/L ammonium acetate solution, and some of the solution underwent LC-MS examination. The analysis was conducted using ultra-high-performance liquid chromatography (UPLC, Ultimate 3000, Thermo Scientific) – electrospray ionization (ESI) – tandem mass spectrometry (MS, Q Exactive Plus, Thermo Scientific), following a previously described method with some modifications [26]. The chromatographic column used was the Hypersil Gold C18 (3μm, 10 mm x 2.1 mm). The mobile phase, consisting of solvent C (50 mM ammonium formate solution) and solvent D (acetonitrile), was delivered at a flow rate of 250 µL/min with the subsequent program: 0–1.00 min, 95% C/5% D; 1.00–1.10 min, a gradual change from 95% C/5% D to 75% C/25% D; 1.10–7.00 min, a gradual change from 75% C/25% D to 60% C/40% D; 7.00 – 8 min, 60% C/40% D. The quantification of CoA-esters was conducted with selected ion monitoring mode in positive polarity. All genuine CoA-ester reference compounds were obtained from Cayman Chemical.

### 2.5. Metabolic network model reconstruction, simulation, and analysis

The reconstruction process strictly adhered to the protocol recommended by Thiele, I [27]. In this study, we incorporated the n-butanol metabolic pathway into the model, which is detailed in supplement.

Constraint-based FBA was conducted using the COBRA toolbox and Gurobi 9.5.2 as the linear programming optimization solver, with metabolic network’s stoichiometric matrix providing the constraints. Linear programming algorithms were employed to compute the optimal solution within the space of feasible flux solutions. During this study, our primary focus was on maximizing FK520 product synthesis rates as the biological phenotype of the optimal solution. This problem is formulated using matrix notation and is presented as Equations (1)–(3) [28,29].

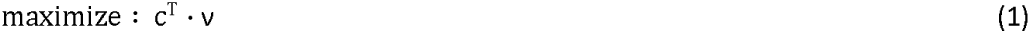

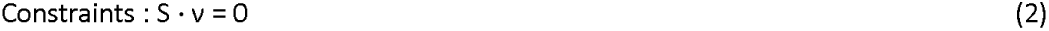

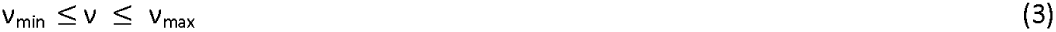

S denotes a stoichiometric matrix encompassing stoichiometric coefficients of metabolites participating in metabolic reactions, while ν represents a vector containing the fluxes of these metabolic reactions. The parameters ν_min_ and ν_max_ correspond to the minimum and maximum constraints, signifying the maximum enzymatic reaction rate and the reversibility of metabolic reactions, respectively. The vector C^T^ signifies a weight vector that suggests each reaction’s contribution on the objective function. Glucose was selected as the only carbon source used in the simulations, all simulations were executed using MATLAB 2023a platform. The principle of robustness analysis was first proposed by Edwards JS [30]. The complete code of robustness analysis can be accessed in Supplementary Material (S1).

### 2.6 Statistical analysis of data

All research data are means of three replicates and all research data are reported as mean ± standard deviation (SD). To evaluate variations across various treatments, we conducted an analysis of variance (ANOVA) using Prism software.

## 3. Results and discussion

### 3.1. short-chain alcohols facilitate FK520 accumulation

Firstly, we attempted to supply five short-chain alcohols (n-propanol, isopropanol, n-butanol, isobutanol, 1,3-propanediol) during the fermentation in *S. hygroscopicus* ATCC 14891 and explore their influence for the production of FK520. As shown in Fig. 1a, the supplement of 1.2% n-propanol, 1.2% n-butanol, and 1.2% isobutanol could significantly stimulate the production of FK520. The highest yield was 375.47 ± 50.35 mg/L with 1.2% n-butanol addition at 24 h, a 2.08-fold enhancement over the control group. And the supplement of 1.2% n-propanol at 24 h also increase the production of FK520 by 1.73-fold. Likewise, 1.2% isobutanol addition at 24 h increase the FK520 production by 1.59-fold. To assess the impact of short-chain alcohols on FK520 titer, we further evaluated various externally added concentrations of n-propanol and n-butanol during FK520 fermentation.

Fig. 1b illustrates that the FK520 production exhibited a certain level of improvement with the content of n-propanol from 0.8% to 1.4% (v/v). All concentrations of n-propanol clearly promoted the overproduction of FK520. Nevertheless, no significant distinctions were observed within experimental groups. The highest FK520 yield was 364.98 ± 41.39 mg/L with 0.8% n-propanol addition.However, the supply of n-propanol also significantly elevates the content of an impurity with a peak very close to that of FK520 (Fig. 2a). As we know, one difference between FK520 and its structural analog such as FK506 lies in the different biosynthesis precursors of C21. The former uses ethylmalonyl-CoA as precursors to build FK520. In previous report, n-propanol could be used to synthesis methylmalonyl-CoA in *Streptomyces erythreus* [19]. Methylmalonyl-CoA competes with ethylmalonyl-CoA for the carbon skeleton at position 21 (Fig. 2c), resulting in the strain producing a structurally similar compound, FK523, alongside the synthesis of FK520 [31]. To confirm whether the impurity produced during fermentation under the conditions of adding n-propanol is FK523, we conducted UPLC-MS qualitative analysis on the fermentation extract. The molecular formula of FK523 is C_42_H_67_NO_12,_ and the electrospray mass spectrum (Fig. 2d) of the fermentation extract with n-propanol revealed the existence of sodium adducts at m/z 800, with fragment ions detected at m/z 782[(M − H_2_O)Na]^+^, 689 [(M − 111)Na]^+^, 671[(M − 111 − H_2_O)Na]^+^, 627[(M − 111 − H O − CO)Na]^+^, 590 [(M − 210)Na]^+^, 479 [(M − 210 − 111)Na]^+^, based on the analysis of fragment characteristics of FK506 and structurally similar compounds as reported in Lhoest, G’s work [32], we have confirmed that the impurity produced in large quantities upon adding n-propanol to *S. hygroscopicus* ATCC 14891 is a structural analogue of FK520, namely FK523. Therefore, while the addition of n-propanol increases the yield of FK520, the large amounts of close structural resemblance of FK523 (the peak area of FK523 reached 40% of FK520) undoubtedly complicates the later stages of FK520 isolation and extraction.

To further enhance the yield of FK520 with addition of n-butanol, we first optimized the oxygen supply conditions in shake flask fermentation and the relevant results and analysis are described in section 3.4. Then, we conducted experiments with the different concentrations of n-butanol (0.4%, 0.8%, 1.2%, 1.6%) as the experimental groups (Fig. 1c). The data revealed that 0.8% n-butanol supply yielded best results, with a production of 523.57 ± 10.08 mg/L, representing a 1.77-fold rise in comparison to control group. Slightly less favorable results were obtained with 1.2% n-butanol (486.77 ± 23 mg/L), while the addition of 1.6% n-butanol had a pronounced inhibitory effect on FK520 production. Next, to explore the influence of the timing of n-butanol addition on FK520 accumulation, We added 0.8% n-butanol at five different time points throughout the whole fermentation period (Fig. 1d). Results indicated that adding 0.8% n-butanol at 27 h yielded the best results with a production of 569.37 ± 24.07 mg/L. Early addition of n-butanol (at 18 h or 21 h) during fermentation significantly inhibited FK520 production. Similar findings have been reported in Li, M’s work [33], where the early addition of n-propanol in Streptomyces natalensis has also led to suppression in mycelia growth and a decrease in Natamycin production. To further investigate the reasons behind the enhanced FK520 production due to n-butanol addition, we conducted additional monitoring and analysis of fermentation process parameters with 0.8% n-butanol supplementation in subsequent experiments.

**Fig. 1.**
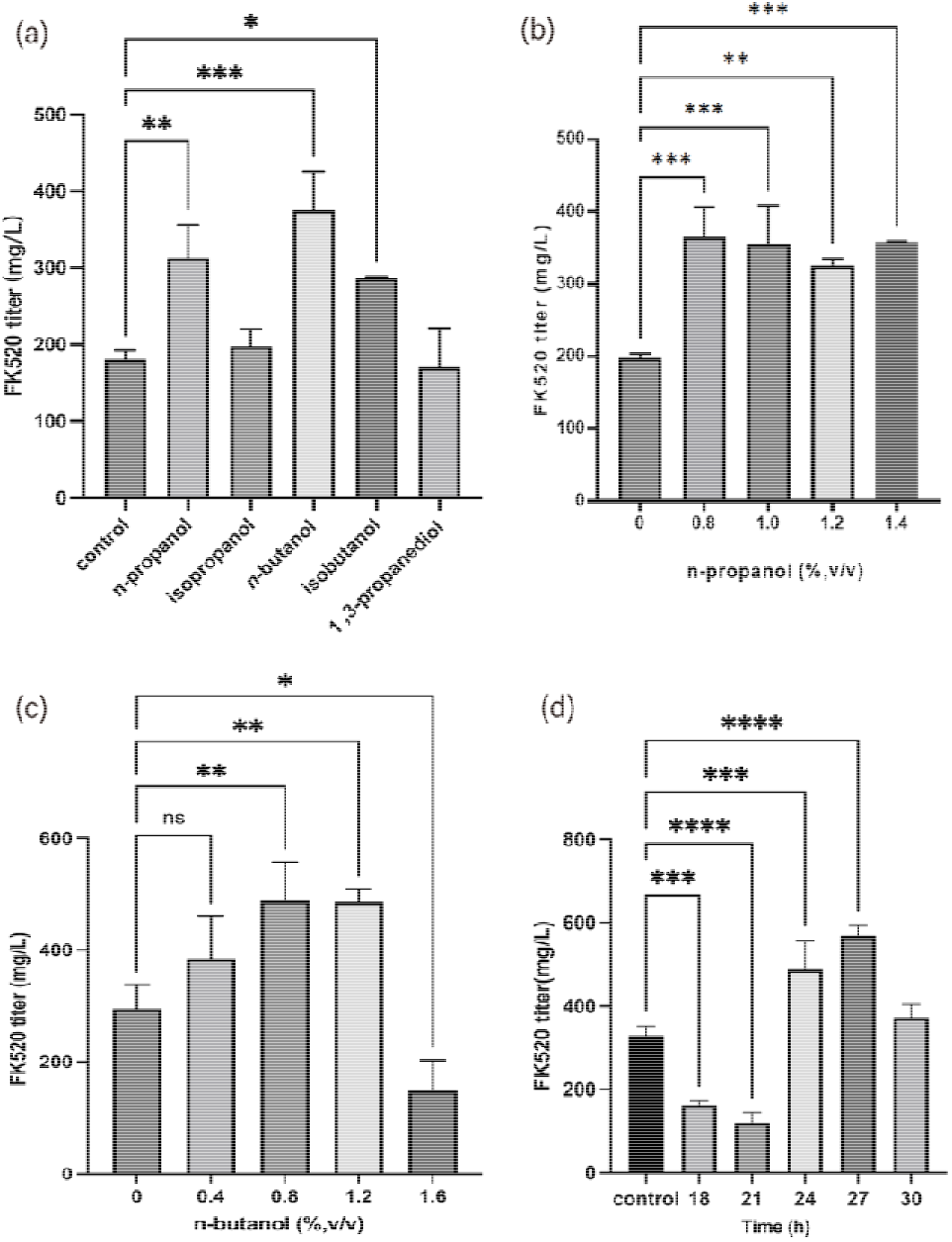
Effects of various short-alcohols on FK520 titer in *S. hygroscopicus var. Ascomyceticus* ATCC 14891. (a) Impacts of supplementing with n-propanol, isopropanol, n-butanol, isobutanol, and 1,3-propanediol on the production of FK520. (b) Investigation of the influence of varying n-propanol concentrations on FK520 production. (c) Investigation of the influence of varying n-butanol concentrations on FK520 production. (d) Impacts of varying time points of addition on FK520 production with 0.8% n-butanol supply. p-values were computed through one-way ANOVA for the respective means (*p <ll0.05, **p < 0.01, ***p <ll0.001, ****p<0.0001) Note: fermentation for (a) and (b) was carried out at a rotational speed of 220 r/min, while fermentation for (c) and (d) was conducted at a rotational speed of 250 r/min.

**Fig. 2.**
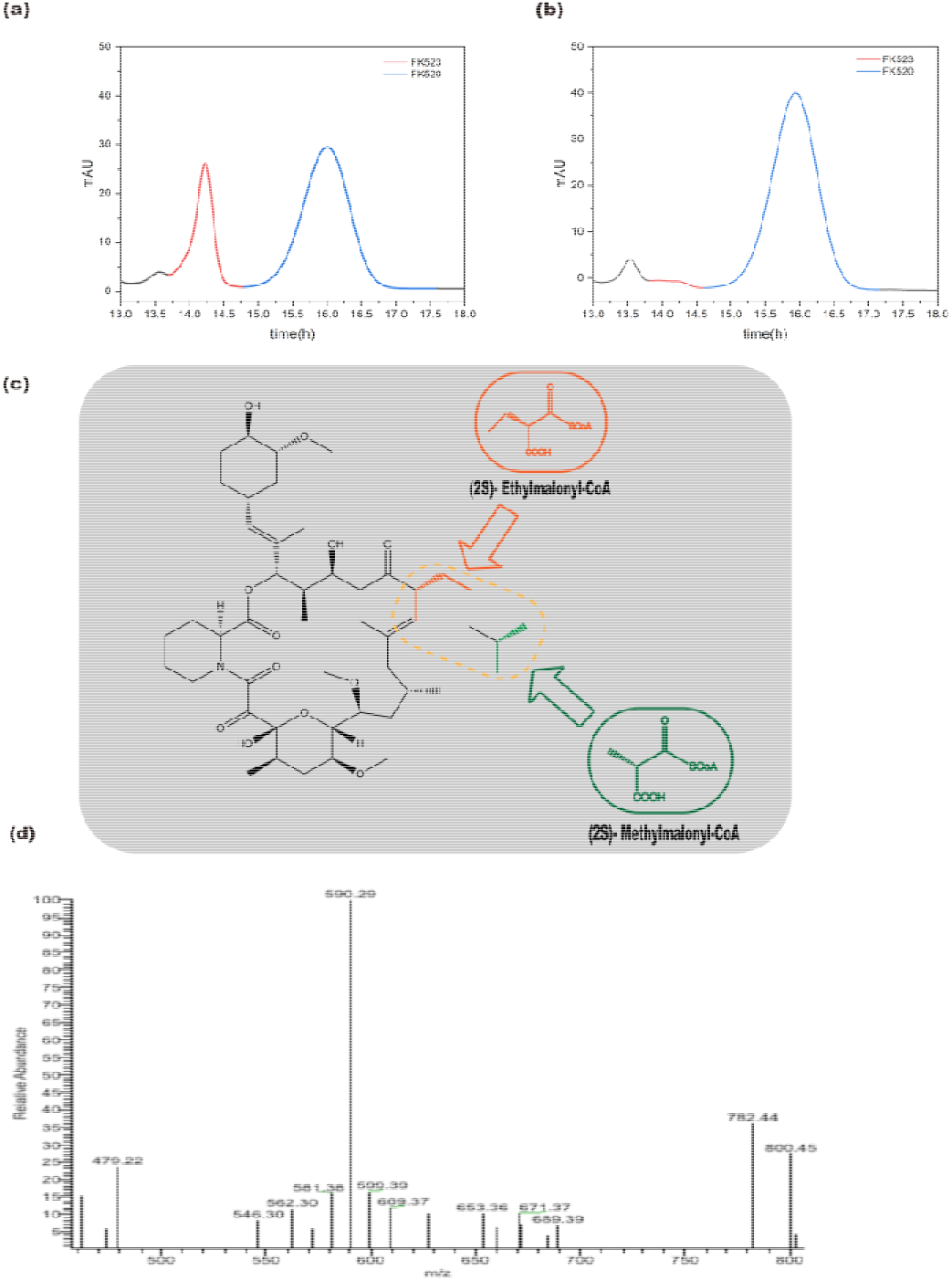
(a) HPLC plots of strains with concentration of 0.8% n-propanol which was added at 24h. (b) The HPLC plots of strains with concentration of 0.8% n-butanol which was added at 27h. (c) In the suggested biosynthetic pathway of CoA-esters within FK520 biosynthesis, the ethylmalonyl-CoA component, indicated in orange, constitutes the C21 segment of the FK520 structure. In contrast, the C21 segment of FK523 is produced using methylmalonyl-CoA, as denoted in green. (d) Electrospray mass spectrum of compound FK523.

### 3.2. Effects of n-butanol addition on fermentation process parameters

#### 3.2.1 Influence on the pH, Glycerol, FK520 titer and DCW

As shown in Fig. 3a, glycerol consumption exhibited significant changes between the groups with and without n-butanol. For the strains fermentation with n-butanol supplementation, the glycerol content at 96 h and 120 h markedly exceeded those of the control group. This indicates that n-butanol supply altered the strain’s utilization of carbon sources especially at the late stage of FK520 fermentation. One reason for the decreased glycerol utilization capacity of *S. hygroscopicus var. ascomyceticus* ATCC 14891 under n-butanol addition conditions may be that GAPDH activity was inhibited with presence of short-chain alcohols [20]. The short-chain alcohols supply may also lead to a rise in Triacylglycerols (TAGs) content [34], and the storage of TAGs is a pivotal factor in controlling the flow of carbon towards the biosynthesis of secondary metabolites, while concurrently decreasing the carbon flow into TCA cycle [35].

In the early stages of fermentation (0-24 h), the pH slowly increased, followed by a rapid decrease over a short period (24-48 h), and then stabilized. When 0.8% n-butanol was added at 27 h, we observed a significant slowdown in pH reduction, and throughout the subsequent fermentation process, the pH level consistently exceeded that of the control group, indicating that 0.8% n-butanol addition inhibited the glycolytic pathway and led to a decrease acid-producing capacity in *S. hygroscopicus var. ascomyceticus* ATCC 14891 and caused a higher pH compared with the control group after 0.8% n-butanol addition. Additionally, in the group with 0.8% n-butanol supply, pH was consistently held around 5.6 on the 3rd, 4th, and 5th days of fermentation. Previous research by Liu has demonstrated that maintaining a pH of around 5.6 in a 5L bioreactor is highly conducive to the accumulation of FK520 [36]. We suppose that the appropriate environmental pH could promote the biosynthesis of FK520, leading to the observed significant accumulation of FK520 in the late stage of fermentation. Current perspectives suggest that antibiotics may function as ‘collective regulators of microbial community homeostasis’ functioning as signals instead of weapons [37,38].Hence, in response to the pressure exerted by the 0.8% n-butanol supply, cells may adjust the environmental pH to promote the biosynthesis of FK520. and use FK520 as a signaling molecule to promote symbiotic relationships among different organisms, benefiting each organism through nutrition or protection.

Despite the reduced glycerol utilization rate, it was evident from the Dry Cell Weight (DCW) data that the supply of n-butanol did not result in a notable reduction in biomass. This indicated that *S. hygroscopicus var. ascomyceticus* ATCC 14891 exhibited a certain degree of tolerance to n-butanol and could ease toxicity by altering its metabolism. According to the literature by Halan B[16], under conditions with a high concentration of n-butanol, medium supplementation with yeast extracts and other substances can alleviate toxicity by lowering the expenses associated with supporting biofilm formation via direct provision of precursors, such as amino acids. Therefore, we speculate that the appropriate addition of n-butanol promotes the utilization of amino acids and other precursors and then redirects more carbon flux towards the biosynthesis of secondary metabolites in *S. hygroscopicus var. ascomyceticus* ATCC 14891. As fermentation progresses from 0 to 120 h, the production of FK520 gradually increases. However, we observed that the synthesis rate of FK520 lagged behind that of control group at 48 h. We supposed that this could be attributed to the addition of 0.8% n-butanol at 27 h, which imposed a certain level of stress on the strain’s growth and metabolism, disrupting its normal growth and metabolic activities. Consequently, this led to a temporary reduction in the synthesis rate of FK520. Subsequently, as time progressed, the strain adapted to the metabolic stress induced by the supply of n-butanol and began to produce FK520 in larger quantities.

In summary, we conclude that supply of 0.8% n-butanol at 27 h alters the internal metabolism of the strain and promotes the biosynthesis and accumulation of FK520. To explore the reasons behind the metabolic changes induced by 0.8% n-butanol addition, we conducted further exploration and analysis of other possible changes during the fermentation period.

**Fig. 3.**
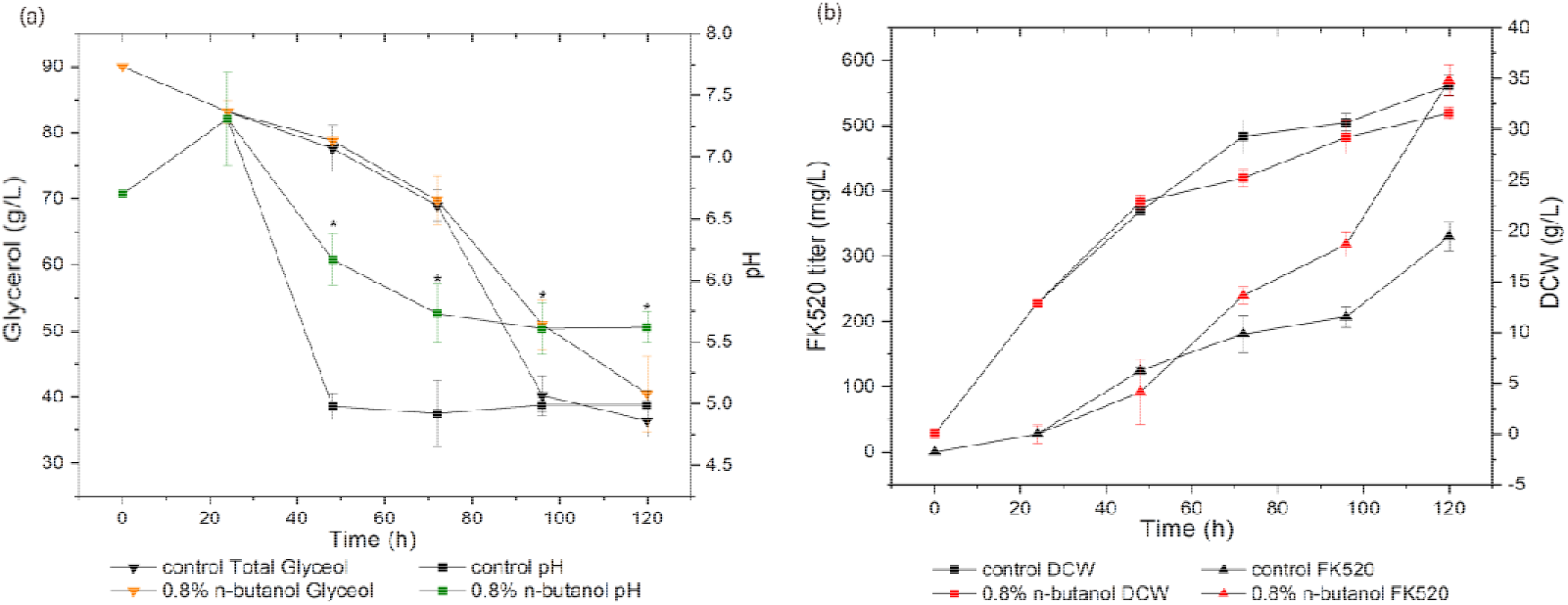
Influences of n-butanol supply on FK520 biosynthesis in *S. hygroscopicus var. Ascomyceticus* ATCC 14891. (a) The fermentation profiles of pH and glycerol following the supply of 0.8% n-butanol at 27 h. (b) The fermentation profiles of biomass and FK520 following the supply of 0.8% n-butanol at 27 h, p-values were computed through one-way ANOVA for the respective means (*p <ll0.05).

#### 3.2.2 Effects of exogenous n-butanol stress to metabolism

n-butanol exhibits significant solubility within cellular membranes, with a maximum concentration of 1.59 M, leading to its incorporation into the membrane. This incorporation interrupts hydrogen bonds among lipid tails, leading to changes of the cell membrane [39].

Furthermore, according to literature reports in Rutherford, BJ’s work [40], oxidative stress is one of the most potent stress responses induced by the addition of n-butanol in Escherichia coli. In this experiment, we sampled at 2 h and 9 h after the addition of n-butanol to assess intracellular ROS changes. The results showed that after adding 0.8% n-butanol at 27 h of fermentation, intracellular ROS levels rapidly increased within 2 h, reaching 1.49x10 RFU/g biomass at 29 h, which is 3.02 times higher than the control. When ROS levels increased, antioxidant proteins responsible for oxidative stress responses, including PerR, SoxRS, RpoS, and OxyR, coordinate a cascade of reactions to mitigate the excessive ROS levels within the bacterial cells. [41,42,43,44]. Therefore, we observed that the ROS levels at 36 h, while still higher than the control group, had significantly decreased compared to 29 h, indicating continuous elimination of ROS through bacterial decomposition and metabolism.

n-butanol-induced ROS also serves as a signaling molecule and play a significant part in modulating both cell growth and secondary metabolism. [45,46]. Previous studies have indicated that ROS can upregulate key genes responsible for enzymes involved in substrates like acetyl-CoA and malonyl-CoA through the oxidative stress response. [47,48], and accumulation of acetyl-CoA and malonyl-CoA during fermentation process was also been observed (Fig. 5c and 5d). We suppose that this accumulation may be related to the accumulation of ROS. Furthermore, research suggests that PHB biosynthesis may be enhanced by ROS-induced stress [49], and PHB could degraded by FkbU to 3-hydroxybutyryl-CoA, and then further degraded by FkbE to ethylmalonyl-CoA in Streptomyces hygroscopicus ATCC 14891 [50,51]. Additionally, ROS can oxidize unsaturated fatty acids, leading to the formation of aldehydes such as malondialdehyde, which can damage proteins and DNA [52] and then cause cellular damage or even cell lysis and death. This may also be one of the reasons for the decrease in FK520 production with a higher n-butanol concentration (1.6%) (Fig. 1c).

Therefore, as n-butanol supplement during the fermentation, the n-butanol or the n-butanol-induced ROS could damage cell membranes. A measure that bacteria respond to environmental stress and membrane damage is altering membrane composition, such as synthesizing more saturated or unsaturated fatty acids [53,54]. As a precursor of intracellular fatty acids, in our opinion, the accumulation of malonyl-CoA (Fig. 5d) after the addition of n-butanol were related to changes in intracellular fatty acid composition aroused by the n-butanol or the n-butanol-induced ROS.

**Fig. 4.**
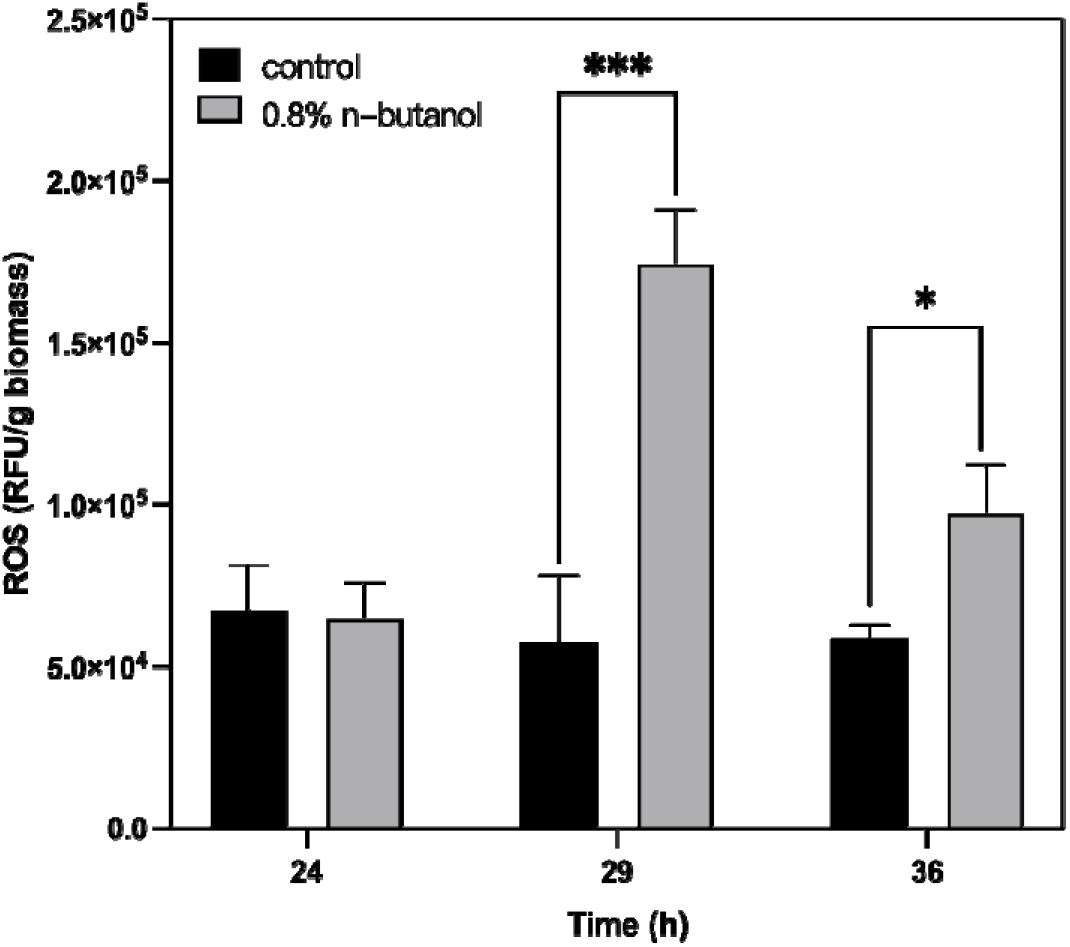
Time course of ROS changing when 0.8% n-butanol were supplied at 27h. p-values were computed through one-way ANOVA for the respective means (*p <ll0.05, ***p <ll0.001).

### 3.3 Intracellular CoA-esters change aroused by n-butanol

Apart from its regulatory role, we hypothesize that n-butanol may also participate in the metabolic reactions of the bacteria by promoting the accumulation of direct precursors of FK520 biosynthesis. To investigate this, we monitored the levels of several important FK520 biosynthesis precursors (methylmalonyl-CoA, ethylmalonyl-CoA, malonyl-CoA, and acetyl-CoA) within S.hygroscopicus ATCC 14891 at different time during fermentation using UPLC-MS. As shown in Fig. 5, when 0.8% n-butanol was added at 27h, the content of ethylmalonyl-CoA rapidly increased, and reached the maximum level at 48h. The result suggests that n-butanol is metabolized within the bacterium to generate butyryl-CoA, which is subsequently converted into the FK520 synthesis precursor ethylmalonyl-CoA, thus facilitating FK520 biosynthesis. Then the ethylmalonyl-CoA was decrease in the 3rd day and we consider that as bacterial metabolism progresses, the supplemented ethylmalonyl-CoA was participating in bacterial metabolic reactions and caused the decrease of itself. Beside of participating in biosynthesis of FK520, ethylmalonyl-CoA can also undergo a series of reactions to be transformed into methylmalonyl-CoA (Fig. 6). Methylmalonyl-CoA serves as both a precursor for FK520 synthesis and can be further converted to succinyl-CoA, entering the TCA cycle, thereby providing energy for bacterial growth. The fluctuation in the content of ethylmalonyl-CoA, with a decrease on the 3rd day followed by an increase on the 4th day, suggested that the intracellular levels of ethylmalonyl-CoA exhibit dynamic and fluctuating patterns. In contrast to the n-butanol supplementation group, the control group exhibited stable content of ethylmalonyl-CoA and methylmalonyl-CoA. Notably, the average concentration of methylmalonyl-CoA in control group was about five times higher than that of ethylmalonyl-CoA. It is known that FK520 biosynthesis necessitates 1mol of precursor ethylmalonyl-CoA and 5mol of methylmalonyl-CoA. The consistent content of ethylmalonyl-CoA and methylmalonyl-CoA in control group suggested that the bacteria actively regulated the proportions of FK520 biosynthesis precursors. As the precursors of FK520 biosynthesis, the higher accumulation of ethylmalonyl-CoA, methylmalonyl-CoA, and malonyl-CoA in n-butanol addition group led to significant increment of FK520 production. Furthermore, the increment of ethylmalonyl-CoA could effectively reduce impurity FK523 content [31]. Therefore, we consider n-butanol addition being exogenous precursor can better promote FK520 biosynthesis than n-propanol addition.

In comparison to increasing the flux of ethylmalonyl-CoA synthesis pathway through genetic engineering [31], the addition of n-butanol also promoted the biosynthesis of methylmalonyl-CoA and malonyl-CoA (Fig.5b and 5d), which are not observed in Yu’s research. This suggests that, apart from serving as a precursor supplement for ethylmalonyl-CoA, the addition of n-butanol also regulates the intracellular carbon flux towards the accumulation of other precursors in the FK520 biosynthesis. After the supply of isobutanol in *A.pretiosum*, a significant increase in malonyl-CoA content was also been detected [18], so we suppose that short-chain alcohols supplement may promote the accumulation of malonyl-CoA in Actinomycetes. As an extension unit for the carbon skeleton, malonyl-CoA plays a significant part in FK520 biosynthesis. The changing trend of malonyl-CoA content closely mirrors the intracellular content of FK520 (Fig. 3b). Furthermore, there is a rapid increase in malonyl-CoA content from the 4th day to the 5th day, coinciding with the period of significant FK520 accumulation. Yu’s work demonstrates that through genetic engineering means, increasing the malonyl-CoA content can enhance FK520 production by 73.3% [31], which means that malonyl-CoA also play important roles in biosynthesis of FK520. Acetyl-CoA is an curcial precursor metabolite that plays a central part in the synthesis of lipids,amino acids, polyketides, isoprenoids, and a wide range of other bioproducts. As a central node molecule, its levels are tightly regulated by the cell’s central metabolism [55]. Despite its high robustness compared to other CoA-esters, we did not observe significant changes in response to n-butanol addition. However, starting from the 3rd day, the average acetyl-CoA content was consistently greater than that of the control group, suggesting that the supply of n-butanol may have a discernible impact on acetyl-CoA accumulation.

**Fig. 5.**
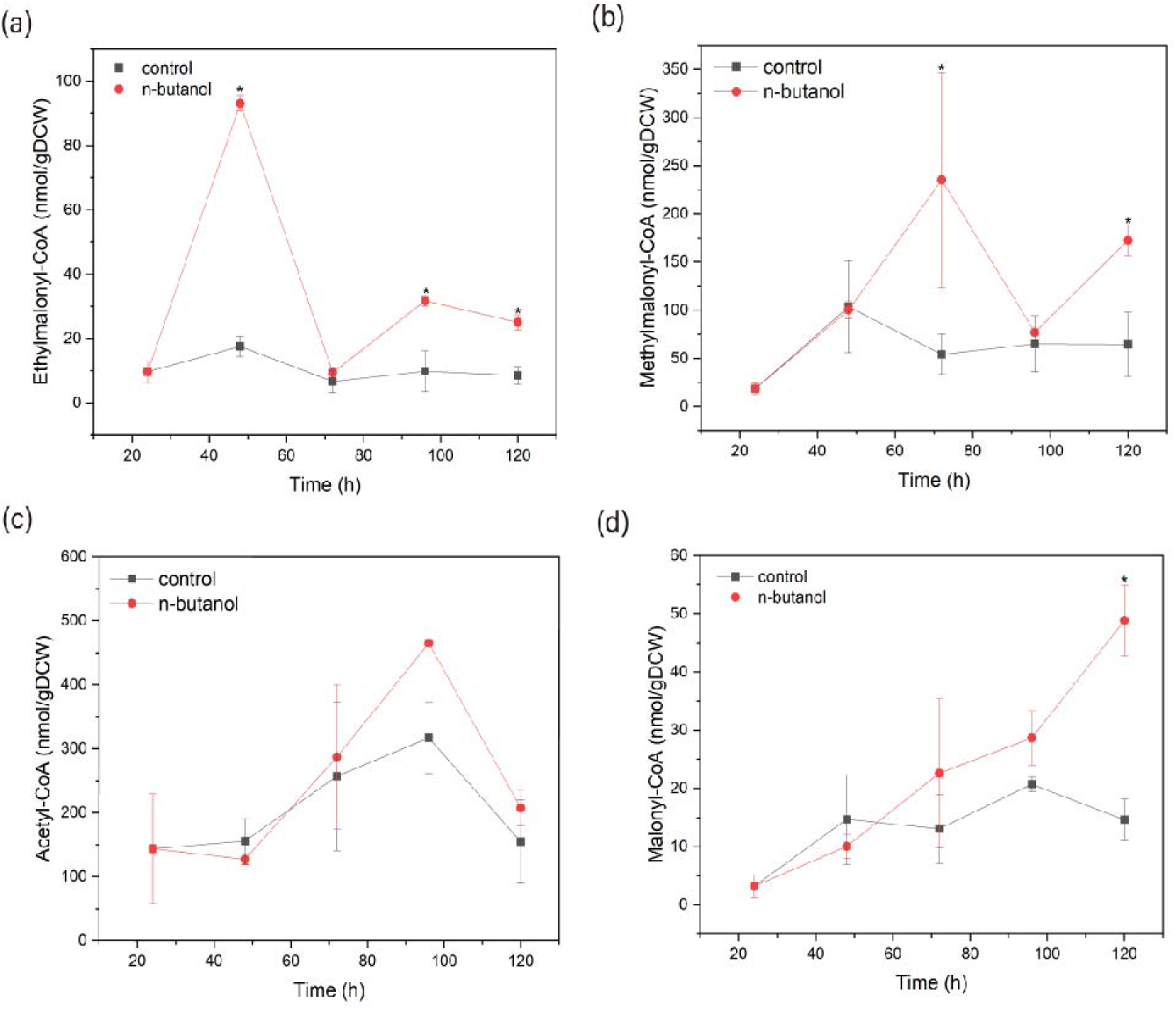
Time course of concentrations of intracellular CoA-esters in *S. hygroscopicus var. ascomyceticus* ATCC 14891 supplemented with 0.8% n-butanol addition (a) Ethylmalonyl-CoA. (b) Methylmalonyl-CoA. (c) Acetyl-CoA. (d) Malonyl-CoA. p-values were computed through one-way ANOVA for the respective means (*p <ll0.05).

### 3.4 Genome-Scale Metabolic Modeling (GSMM) application of Optimization for n-butanol Supplementation

#### 3.4.1 Analysis of n-butanol addition on FK520 metabolism

**Fig. 6.**
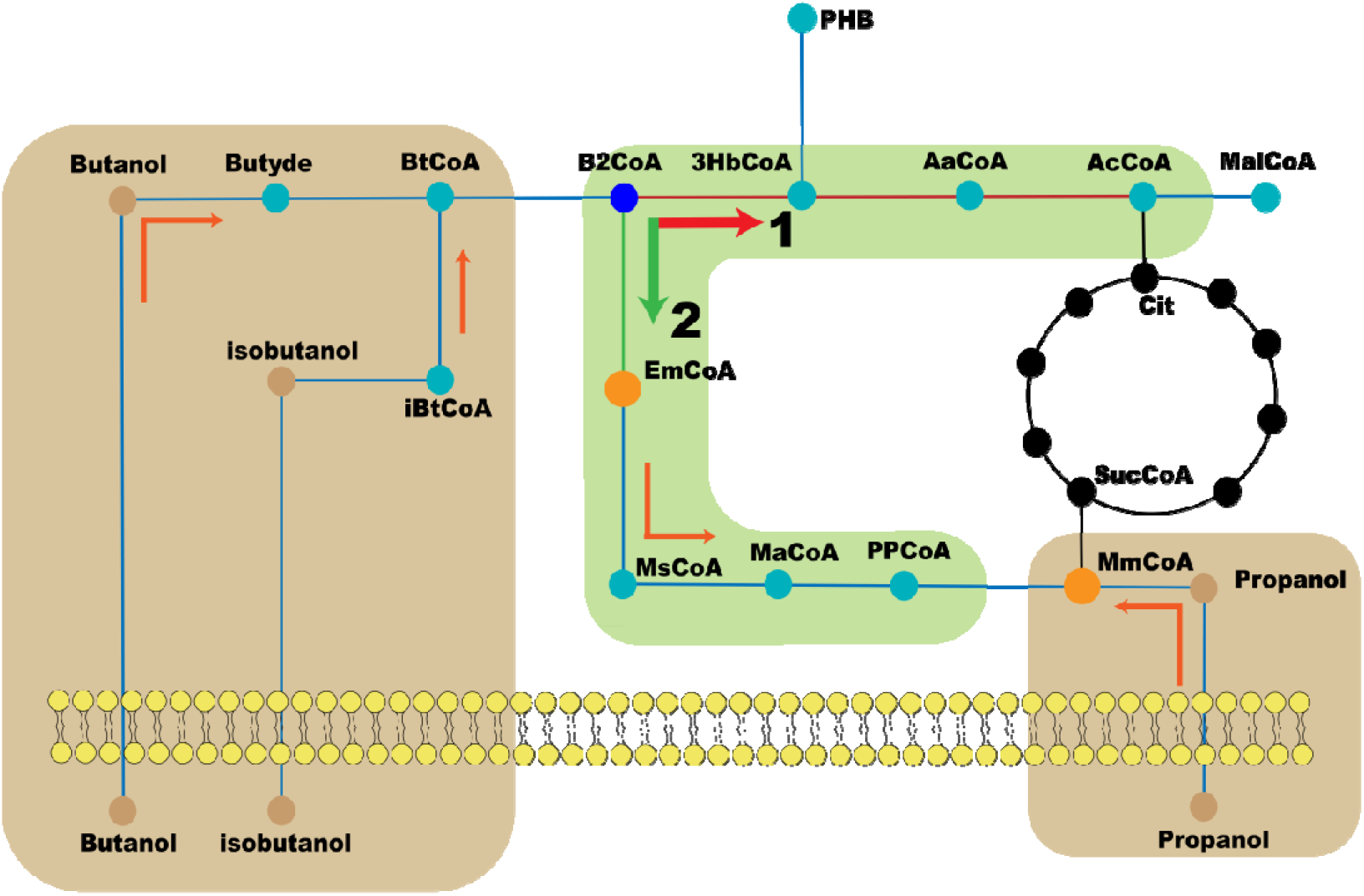
Main alterations in *S. hygroscopicus var. ascomyceticus* response pathways with short-chain alcohols treatment adding n-proponal, n-butanol, and isobutanol to the cultivation medium (brown); catabolic pathway of crotonoyl-CoA (green). Butyde, butyraldehyde; BtCoA, butyryl-CoA; B2CoA, Crotnnoyl-CoA; 3HbCoA, 3-hydroxybutyryl-CoA; AaCoA, acetoacetyl-CoA; cit, citric acid; EmCoA, (2S)-ethylmalonyl-CoA; MsCoA, methylsuccinyl-CoA; MaCoA, (2R,3S)-β-methylmalyl-CoA; PPCoA, propionyl-CoA; MmCoA, (2S)-methylmalonyl-CoA; SucCoA, succinylCoA; MalCoA, malonyl-CoA;

To further analyze the impact of n-butanol addition on the metabolic network of *S. hygroscopicus var. ascomyceticus* ATCC 14891, we utilized a previously reported GSMM (the maximal specific growth rate was 0.0653/h) [56] and incorporated n-butanol matabolic reaction tostimulate and analyze the metabolism of *S. hygroscopicus var. ascomyceticus* with n-butanol addition. In Streptomyces coelicolor A3(2), n-butanol were converted to Butyraldehyde through alcohol dehydrogenase and then further transformed into butanoyl-CoA under the action of acetaldehyde dehydrogenase [57]. Because of the high conservation of enzymes in different Streptomyces species [56], we could refer to the metabolic pathway of n-butanol in Streptomyces coelicolor A3(2), and reactions were been incorporated into the model.

The robustness of n-butanol supplementation was analyzed on the specific FK520 synthesis rate using GSMM. The results indicate that when the n-butanol addition rate is 0 mmol/gDCW/h, the shadow price of n-butanol is 0.0181 (Fig. 7a). This suggests that an increase of 1 mmol/gDCW/h in the n-butanol uptake rate will lead to an increase of 0.018 mmol/gDCW/h in FK520 synthesis rate. As the n-butanol uptake rate increases, the relative FK520 synthesis rate continuously rises (Fig. 7a). Notably, the shadow price of n-butanol are changing with n-butanol uptake rate change, and when n-butanol uptake rate is 2 mmol/gDCW/h and 10 mmol/gDCW/h, the shadow price of n-butanol is 0.0159 and 0.0136, respectively. This demonstrates that n-butanol supplementation, as a precursor for FK520 synthesis, effectively promotes FK520 production. Furthermore, oxygen supply is another key factor of FK520 biosynthesis, and oxygen demand for FK520 biosynthesis was vary in different conditions [58,59]. Therefore, to analyze the impact of oxygen uptake rate on FK520 biosynthesis with n-butanol addition, we employed phenotype phase plane analysis to investigate the interaction between these two parameters and their effect on FK520 specific synthesis rate [60]. For detailed execution code, please refer to Supplementary Material (S2). From Fig. 7b, it can be observed that this surface plot contains two distinct plane, which are separated by segment AB, and both of them are affected by oxygen uptake rate and n-butanol uptake rate. Furthermore, when one parameter is fixed, the other parameter has an optimal value at which qFK520 is maximized, and this point lies on line segment AB. Analysis reveals that all points in plane I have oxygen uptake rates lower than their optimal uptake rates, while n-butanol uptake rates are higher than their optimal rates. Analysis of the flux distribution simulated by GSMM helps us understand the features of plane I. When oxygen supply is insufficient, the excessive n-butanol addition will lead to insufficient carbon sources to maintain the desired biomass. At this point, a large amount of n-butanol is converted into actyl-CoA and enters the TCA cycle (Pathway 1 in Fig. 6) as the second carbon source to support cell growth. In this condition, reducing the n-butanol uptake rate and increasing oxygen uptake rate allow more n-butanol to flow towards the biosynthesis of FK520 precursors (pathway 2 in Fig. 6), thereby facilitating the biosynthesis of FK520. In contrast, the characteristics of plane II are the opposite of plane I, which all points have oxygen uptake rates higher than their optimal uptake rates, while n-butanol uptake rates are lower than their optimal rates. Due to the inability of fed-batch culture in shake flask fermentation, We suppose that the poor oxygen supply capacity of flask fermentation may limit the organism’s ability to utilize n-butanol for FK520 biosynthesis, and increasing oxygen supple could ease that limitation.

**Fig. 7.**
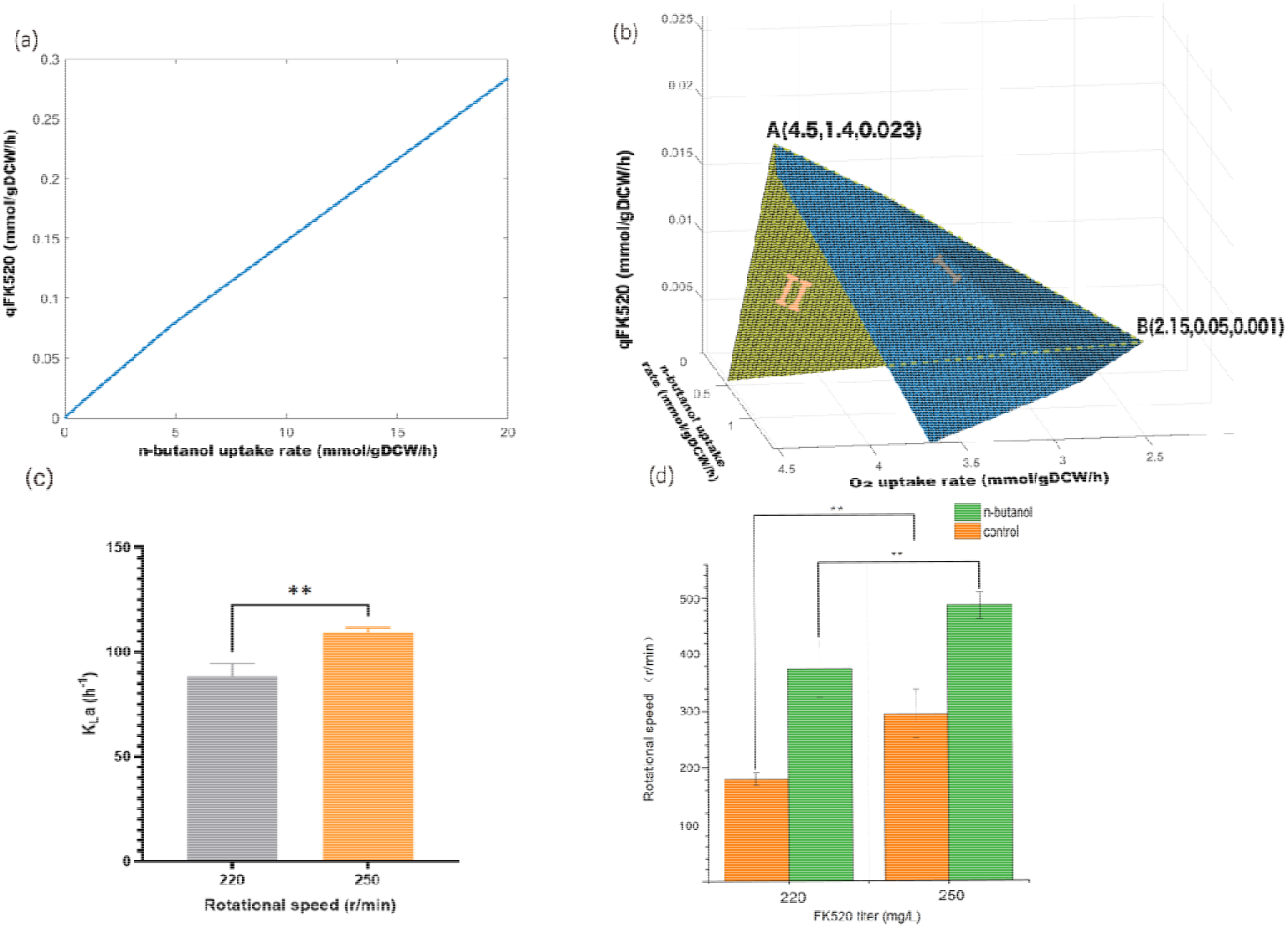
(a) Robustness analysis of the n-butanol uptake rate to the specific FK520 synthesis rate with infinity oxygen. (b) Phenotypic phase planes analysis for FK520 synthesis rate with varying n-butanol and oxygen uptake rates. (c) KLa value during different rotational speed. (d) FK520 titer in different rotational speed with and without 1.2% n-butanol. p-values were computed through Student’s t-test for respective means (**p <ll0.01).

#### 3.4.2 Effects of different rotational speed on FK520 in shake flask fermentation

In flask fermentation, common methods to enhance oxygen supply include using baffled shake flasks and increasing the shaking speed. However, we found that the use of baffled flasks could lead to splashing of the fermentation liquid and greater evaporation. Therefore, we opted to increase the rotational speed to improve oxygen supply capacity in flask fermentation. We raised the rotational speed from 220 r/min to 250 r/min and gauged the KLa of the flask to assess oxygen supplement of flasks [23]. As shown in Fig. 7c and Fig. 7d, when we increased the rotational speed to 250 r/min, the KLa value of the shake flask reached 109.5 h, representing a 24% increase compared to the control. Under this condition, we observed significant increases in FK520 production both with and without the supply of n-butanol, which confirm the results of stimulation. In the group without n-butanol addition, FK520 production reached 295.2 mg/L, a 63.2% increase compared with the oxygen-limited group. In the group with 1.2% n-butanol addition, FK520 production reached 486.77 mg/L, a 29.6% increase compared to the oxygen-limited group. Furthermore, our simulations suggest that, besides change oxygen supply during the fermentation process, controlling the feed rate of n-butanol is also important to promote FK520 biosynthesis. However, controlling the feed rate of n-butanol in flask fermentation presents challenges, so this method requires further investigation in subsequent bioreactor fermentation.

## 4. Conclusion

The supply of 0.8% n-butanol regulates carbon flux within the *S. hygroscopicus var. ascomyceticus* ATCC 14891 towards secondary metabolic reactions, significantly promoting the accumulation of FK520. Besides that, it also directly contributes to the significant supplementation of three key precursor, including ethylmalonyl-CoA, methylmalonyl-CoA, and malonyl-CoA, for FK520 biosynthesis. Furthermore, we found that the increasing oxygen supply could improve the FK520 production in shake flask fermentation with n-butanol addition. Our work provides valuable insights for optimizing the fermentation process of FK520 in the future.

## Supporting information

GSMM

## Acknowledgements

This work was supported by National Key Research and Development Program of China(2020YFA0907800).

## CRediT authorship contribution statement

JM: Conceptualization, Methodology, Formal analysis, Visualization, Investigation, Data curation, Software, Writing – Original Draft. QC: Investigation; YL and DZ and SG: Supervision, Fund acquisition, Writing – Review & Editing. All authors read and approved the final manuscript.

## Data availability

Data will be made available on request.

