## supplemental file1 for "Boosting ascomycin production through n-butanol addition in the fermentation process of *Streptomyces hygroscopicus var. ascomyceticus* ATCC 14891"

model = changeObjective(model,'FK7')

model = changeRxnBounds(model,'EX_Biomass',0.0653,'b')

growthRates=zeros(2000,1);

for i = 0:2000

model=changeRxnBounds(model,'EX_1buol',-i/100,'b')

FBAsolution=optimizeCbModel(model,'max');

growthRates(i+1)=FBAsolution.f;

end

x=[0:0.01:20];

plot(x,growthRates)

S1

model = changeObjective(model,'FK7')

model = changeRxnBounds(model,'EX_Biomass',0.0653,'b')

growthRates=zeros(150,180);

for i=0:150

      model=changeRxnBounds(model,'EX_1buol',-i/100,'b');

    for j=0:180

        model=changeRxnBounds(model,'EX_o2(e)',-j/40,'b');

        FBAsolution=optimizeCbModel(model,'max');

        growthRates(i+1,j+1)=FBAsolution.f;

    end

end

x=[0:0.025:4.5];

y=[0:0.01:1.5];

surfl(x,y,growthRates)

S2
